# Individual Resting-state alpha peak frequency and within-trial changes in alpha peak frequency both predict visual flash segregation performance

**DOI:** 10.1101/2020.05.11.089771

**Authors:** Jan Drewes, Evelyn Muschter, Weina Zhu, David Melcher

## Abstract

Although sensory input is continuous, information must be combined over time to guide action and cognition, leading to the proposal of temporal sampling windows. A number of studies have suggested that a 10Hz sampling window might be involved in the “frame rate” of visual processing. To investigate this, we tested the ability of participants to localize and enumerate 1 or 2 visual flashes presented either at near-threshold (NT) or full contrast intensities, while recording magnetoencephalography (MEG). Performance was linked to the alpha frequency both at the individual level and trial-by-trial. Participants with a higher resting state alpha peak frequency showed the greatest improvement in performance as a function of ISI within the 100ms time window, while those with slower alpha improved more when ISI exceeded 100ms. On each trial, correct enumeration (1 vs. 2) performance was paired with faster pre-stimulus instantaneous alpha frequency. The effect of the timing of the NT stimulus on the perception was consistent with visual temporal integration windows being embedded in a sampling rhythm. Our results suggest that visual sampling/processing speed, linked to peak alpha frequency, is both an individual trait and varies in a state-dependent manner.

**Significance Statement:** A fundamental question in sensory and cognitive neuroscience is how the brain makes sense of the continuous flow of sensory input, parsing it into meaningful objects and events. The speed of cortical alpha rhythms is hypothesized to predict the temporal resolution of visual perception. We present a magnetoencephalography study investigating whether this temporal resolution is an individual trait or, in contrast, depends on fluctuations in brain state. Our results show that both individual resting state alpha frequency, a relatively stable trait, and trial-by-trial fluctuations of the instantaneous alpha frequency determine temporal segregation performance. These results have important implications for how our moment-by-moment perceptual experience is shaped as well as future intervention strategies to improve visual processing for specific tasks.

## Introduction

Sensory information arrives continuously to the brain, raising the question of how cognitive processing, which takes time, is able to act upon the analog flow of information. It is clear that sensory information is combined over time to support tasks such as visual motion processing, multisensory integration or parsing spoken language, as well as to simply improve the accuracy of our perception of stable features of the world. This suggests that perceptual systems process continuous sensory input in discrete “temporal integration windows” (TIWs: (Pöppel, 1997, 2009; Hasson et al., 2008), which may be embedded in sampling rhythms (VanRullen and Koch, 2003; Dehaene, 2016). Specifically, it has been argued that the alpha rhythm provides a natural time frame for temporal integration (Varela et al., 1981; Gho and Varela, 1988).

A neural measure of TIWs in tactile perception has been estimated in an MEG paradigm by quantifying the influence of a near-threshold (NT) tactile stimulus on the evoked response to a second, full intensity stimulus (Wühle et al., 2010, 2011). Wühle and colleagues reasoned that if the inter-stimulus interval (ISI) between the two stimuli exceeded the temporal integration window, then the evoked response to the full stimulus would no longer be affected by the NT stimulus. Across two studies (Wühle et al., 2010, 2011), they found attenuation of the MEG signal in the primary somatosensory (SI) cortex for ISI values up to 60ms, while the attenuation in secondary somatosensory (SII) cortex occurred even for ISIs of several hundred milliseconds. In other words, the temporal resolution of SI was relatively short (within 60ms), while temporal integration within SII lasted over hundreds of milliseconds This finding was partially replicated in another MEG study which also found temporal integration windows for tactile stimuli in the order of 50-75ms (Yamashiro et al., 2011).

The use of paired NT+full stimulation, as used by Wühle and colleagues in the somatosensory domain, could be used to measure the neural correlates of visual TIWs. Importantly, in terms of links to visual sampling and a potential role for alpha oscillations, a NT stimulus is less likely to reset the oscillatory phase (for a similar logic, see: (Ronconi and Melcher, 2017)). A strong phase reset to the first stimulus can influence processing of the second stimulus (Wutz et al., 2014a), in which case the ISI duration effect might reflect a TIW but not endogenous sampling. It is theoretically useful to distinguish between an oscillation and a series of evoked responses or phase resets (see, for example: (Capilla et al., 2011; Zoefel et al., 2018; Doelling et al., 2019; Lakatos et al., 2019).

The main focus of this study was to investigate the role of sampling rhythms in visual perception by testing whether alpha peak frequency predicted performance in this NT+Full stimulation paradigm. On the one hand, it has been argued that peak alpha frequency (PAF) is an individual trait, with some people showing consistently higher or lower alpha peak frequency across testing sessions (Salinsky et al., 1991). Indeed, a number of studies have linked individual differences in PAF to speed of visual processing. However, recent studies have investigated variation in this peak frequency by measuring rapid alterations in the instantaneous peak alpha frequency (Haegens et al., 2014). Specifically, performance in single trials, in rapid visual tasks, has been found to be related to the instantaneous alpha peak frequency (IAF) on that trial (Wutz et al., 2014b, 2018; Cecere et al., 2015; Samaha and Postle, 2015) with evidence that IAF might be to some degree under top-down control (Wutz et al., 2018).

We measured both individual peak alpha frequency, measured for each individual in separate trials, and instantaneous alpha frequency within each trials. We hypothesized that individuals with a higher PAF would have shorter temporal integration windows and that the instantaneous measure of alpha peak would predict, on single trials, whether participants reported seeing separate flashes or a single fused percept.

## Methods

### Study Design

We tested three main predictions. First, based on the previous studies with tactile stimulation, we predicted that the processing of a full-contrast stimulus would be influenced by a preceding NT stimulus when the inter-stimulus interval was within the TIW (within about 100ms).

Second, we predicted that individual subject performance in the perceptual segregation task should be better for subjects with higher temporal resolution. Under the assumption that the individual resting state alpha peak frequency is an indicator of the speed of a particular subject’s visual system and thereby of the ability to resolve brief temporal events, we hypothesize that those subjects with higher individual alpha peak frequency would improve their performance in earlier ISI values compared to subjects with lower alpha peak frequency. Subjects with high individual alpha frequency should therefore improve mostly in the first interval (33-67ms), while subjects with low individual alpha frequency should improve mainly in the last interval (100-400ms).

Third, based on recent work linking performance in rapid visual tasks with the instantaneous alpha peak frequency (IAF) on that trial (Wutz et al., 2014b, 2018; Cecere et al., 2015; Samaha and Postle, 2015) we hypothesized that we would also find higher IAF on trials in which participants correctly discriminated between 1 and 2 stimuli.

### Experimental Procedure

The task consisted of detecting, localizing and enumerating the number of flashes presented on the display. On each trial, one or two localized visual flashes were presented on a medium (50%) gray background (see Figure 1). Flashes were 2D Gaussian luminance distributions with an approximate total diameter of 1 degree visual field, although the perceived size of the Gaussian blob may have been smaller depending on subject and adjusted contrast. Flashes were shown at two contrast levels, full contrast and near-threshold (NT) contrast. The duration of each flash was set to 8.3ms (one screen refresh, 120Hz). In full contrast flashes, the peak of the blob was white (maximum luminance of the display system), giving maximum luminance contrast against the background. Prior to the main experimental sessions, NT contrast flashes were adjusted by a Quest procedure (Watson and Pelli, 1983) until subjects were just above chance performance (57% correct) at detecting a single threshold flash. To reduce the number of chance hits included in the analysis, flashes (or pairs of flashes) were shown in one of four quadrants, symmetrically arranged around the point of fixation, and randomly chosen for each trial (see Figure 1). Subjects were then asked to identify the quadrant where they had seen the flash, rather than whether they had seen it. Flash eccentricity was 6 degrees from point of fixation. Stimuli were generated on a UNIX computer using Matlab 8.0 (The MathWorks) and the PsychophysicsToolbox Version 3 (Brainard, 1997; Pelli, 1997).

**Figure 1:**
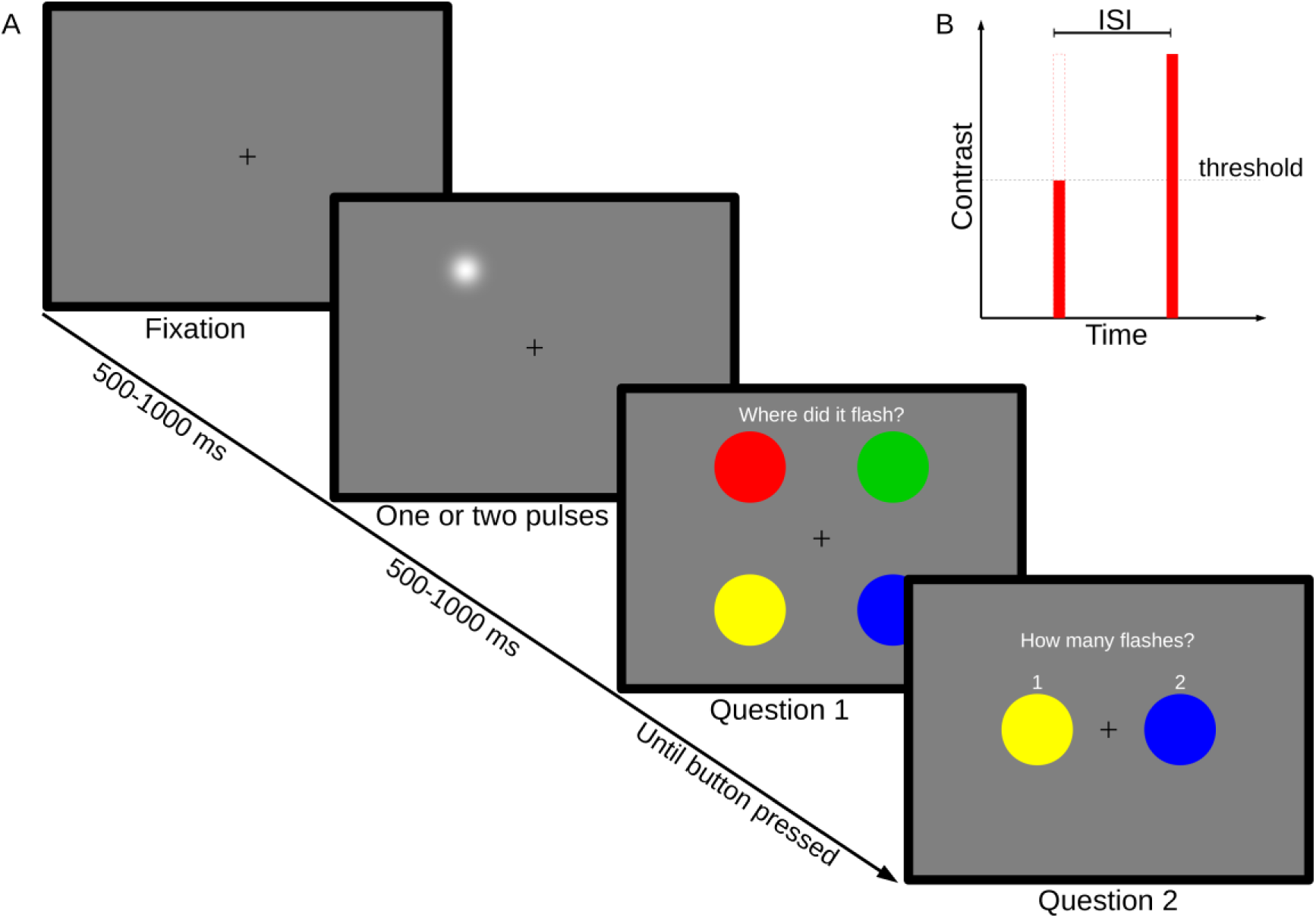
Experimental Paradigm. A) One or two brief flashes of light were shown in one of four quadrants. Subjects pressed colored buttons to indicate their responses. B) Stimulus design. Flashes were either of threshold intensity or full intensity, in the case of dual pulses with a variable ISI.

Each trial was chosen from one of 6 experimental conditions in a randomized fashion. Four dual-flash conditions with different ISIs (33, 67, 100 and 400ms) were intended to probe the temporal integration behavior of the subjects. A single threshold pulse and two full contrast pulses with an ISI of 33ms served as baseline conditions. After each trial, subjects were prompted to make two responses (see Figure 1). The first response was to indicate the screen quadrant in which the flashes were perceived. The second question asked the number of flashes perceived (one or two). Responses were made by means of an MEG-compatible button box, with matching color coding between buttons and displayed questions (see Figure 1). After completion of the main experiment, subjects were directed to look at a fixation cross at the center of the screen, without further tasks. During this time period, 5min of MEG data was recorded for the purpose of resting state analysis.

### MEG Data Acquisition

MEG was recorded during a visual stimulation paradigm. Subjects were seated in an Elekta Neuromag 306 in vertical position, placed inside a magnetically shielded chamber. The display system was a DLP projector (Panasonic PT-D7700E) running at a refresh rate of 120Hz, aimed at a translucent back-projection screen located in a dimly lit, magnetically shielded chamber.

### Participants

Twenty subjects (9 female, mean age 25.6 years, sd=2.39 years, all right-handed) participated in the experiment. All participants provided written informed consent; the study was approved by the Ethical Committee of the University of Trento and was conducted in accordance with the Declaration of Helsinki. Two subjects were excluded from analysis due to magnetic interference, suspected to be due to unreported dental work, and one subject aborted the experiment prematurely.

### Data Processing

#### Behavior/General

Subjects identified the quadrant in which they perceived the visual stimulus; trials on which the wrong quadrant was identified were treated as misses and thus excluded from further analysis (13% of trials, see Results section, below).

#### Event Related Fields

For the trial data analysis, environmental noise was removed and the data was co-registered in order to remove small head movements across the separate measurement runs through signal space separation with spatio-temporal extension (Taulu and Kajola, 2005; Taulu et al., 2005)) implemented via the MaxFilter software version 2.2.15 (Elektra-Neuromag Ltd., Helsinki, Finland). Prior to that, the data was visually inspected and noisy channels were excluded from the tSSS filtering. Data was then analyzed in Matlab using the Fieldtrip toolbox (Oostenveld et al., 2011) for general MEG data treatment and the CoSMoMVPA toolbox for multivariate cluster statistics (Oosterhof et al., 2016). Data was recorded at 1kHz, then downsampled to 250Hz. Epochs of 4s were centered on the stimulus onset (in case of two pulses, on the second pulse). Trials were visually inspected for artifacts and contaminated trials were removed. Of the 1160 trials recorded from each subject, an average of 1016 trials remained.

### Resting State

For the resting state analysis, a ten second long hamming window was moved over the whole 5min resting state recording in 1s steps, and the Fourier amplitude spectra of each window were averaged to form the resting state spectrum of each subject, which was averaged over all channels. The alpha peak of each subject was then identified as the location of the highest amplitude in the alpha frequency range, defined as 7-13Hz. Of the 17 subjects, one subject had to be excluded as resting state data was unavailable due to technical failure. The remaining 16 subjects all had unambiguous alpha peaks (for a sample, see Figure 2) in their resting state data and were entered into the following analysis. On average, individual peak alpha frequency was 10.48Hz (median: 10.6Hz, min: 9.1Hz, max 11.6Hz, see Figure 3). These results are comparable to the values reported in previous studies (Klimesch, 1999; Aurlien et al., 2004; Haegens et al., 2014; Cecere et al., 2015; Wutz et al., 2018).

**Figure 2:**
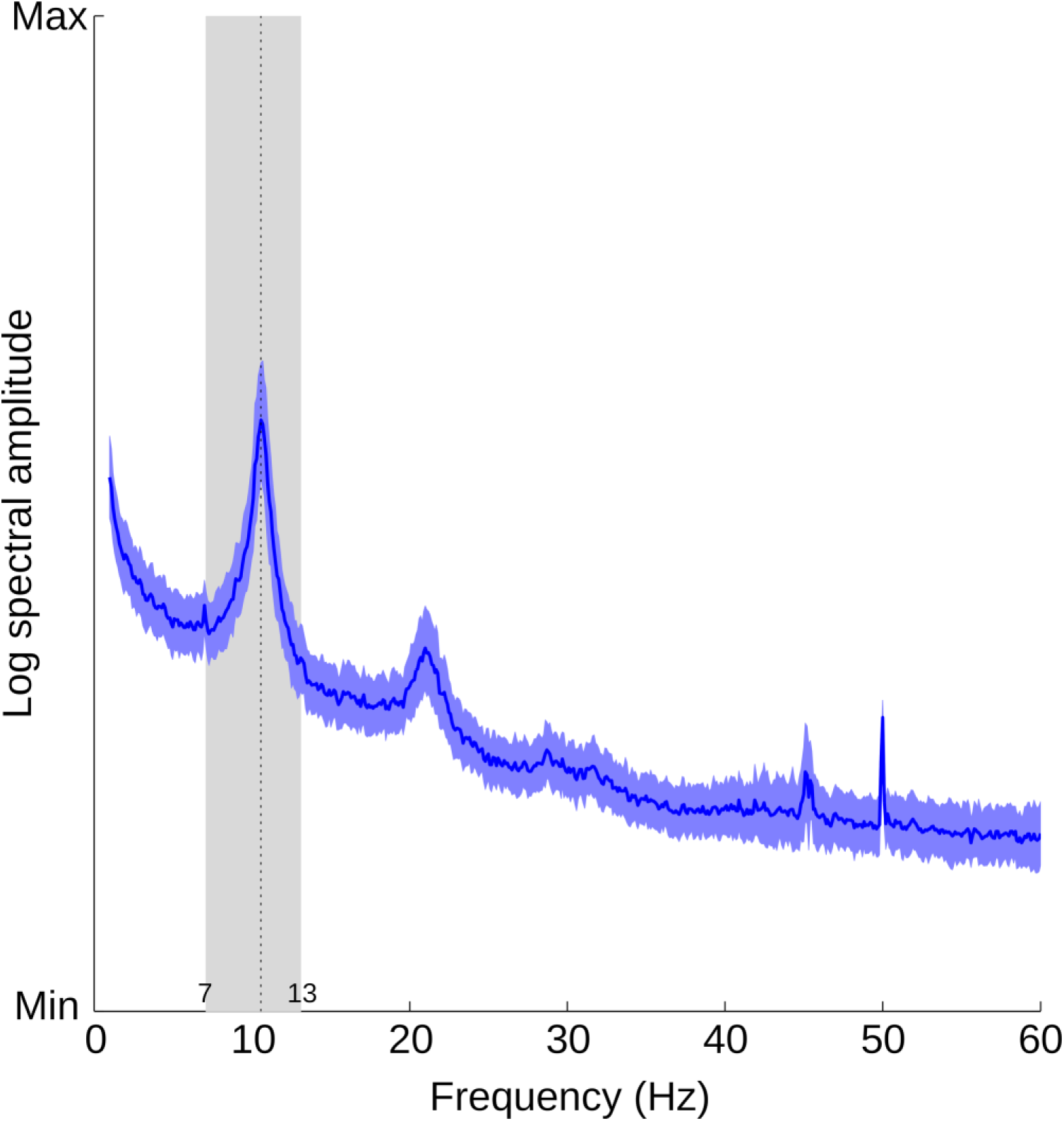
Analysis of individual alpha peak frequency from resting state data. Data from one sample subject. Blue line and shade indicate mean and sd of a 10s sliding window moving over a 5min recording (see Methods). The alpha band (7-13Hz) has been highlighted, and the determined peak in the alpha band marked with a dashed line.

**Figure 3:**
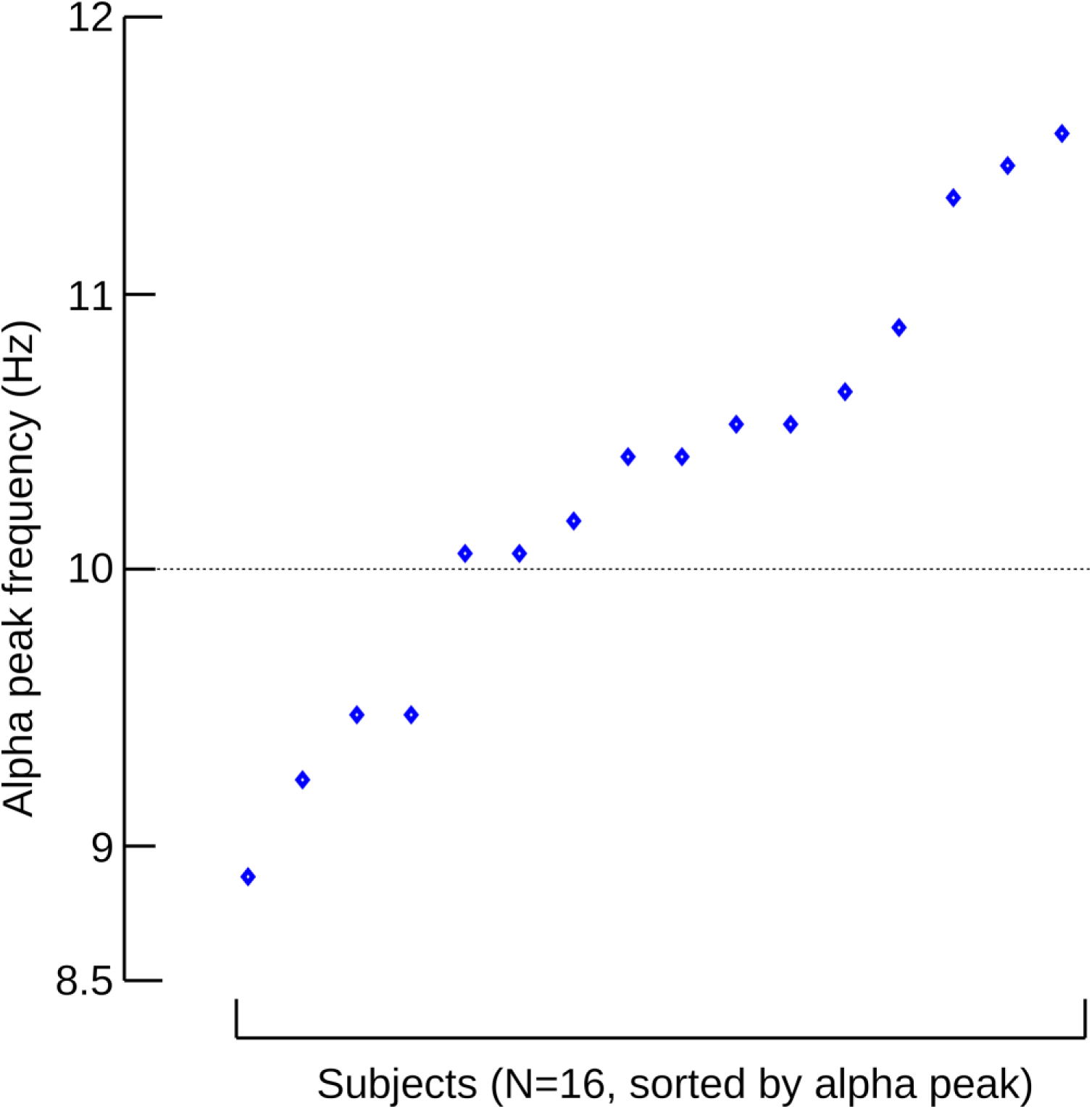
Individual alpha peak frequency for 16 subjects, plotted in ascending order.

### Instantaneous alpha frequency

Experimental data were first band-pass filtered with a zero-phase, plateau-shaped FIR filter in the alpha frequency band between 7 and 13Hz. Then we computed the instantaneous phase angle over time with a Hilbert transformation. As detailed in the publication by (Cohen, 2014), the instantaneous frequency is defined as the time rate of change of the instantaneous phase angle. Thus, the temporal derivate of the instantaneous Hilbert phase corresponds to the instantaneous frequency in Hertz (scaled by sampling rate and 2π). As the resulting phase angle time series is prone to noise that could cause sharp artifacts, we applied a median filter (10 equally spaced window sizes between 10 and 400ms) for ten times. Across those resulting median-filter windows, we calculated the median instantaneous frequency estimates. As task performance was comparable at ISIs of 67ms and 100ms, we pooled this data together and focused our analysis on subjects that reached at least 20% of accuracy (N= 14) in order to obtain adequate statistical power. For each perceptual outcome and for each subject an equal number of trials (in which participants had first correctly identified the correct stimulus location) were randomly selected to prevent any bias across conditions.

We then statistically compared the difference in instantaneous alpha frequency between consciously perceiving one or two flashes. Following the methodology of Samaha and Postle (Samaha and Postle, 2015), we selected the sensors with the highest pre-stimulus power (right occipital: MEG2511 and MEG2541) and then calculated the difference between perceptual outcomes in the pre-stimulus time period from −500ms to 0ms (onset of the 1^st^ flash stimulus) with a dependent samples t-test with correction for multiple comparison by means of an nonparametric cluster-based permutation procedure (Oosterhof et al., 2016).

## Results

### Behavioral Data

On average, subjects detected the stimulus (i. e. identified the correct quadrant) on approximately 87% of all trials (range: 67.2 – 95.8%). Detection performance was distributed homogenously across quadrants (sd of quadrant means 4.0%). The behavioral result after elimination of misses is shown in Figure 4. On average, performance correct (identifying the number of flashes shown) for the NT + Full condition ranged from 22% at 33ms ISI to just less than 50% at 400ms ISI and was therefore within expectation, given that the probability of detecting the threshold pulse was by design adjusted to 57%. Performance on the Full + Full condition was around 34% and therefore higher than the NT + Full condition with the same ISI (33ms). Correct performance with only a single threshold pulse was just below 90%, showing that participants reported seeing only a single flash (rather than two) when the single NT stimulus was presented and correctly localized.

**Figure 4:**
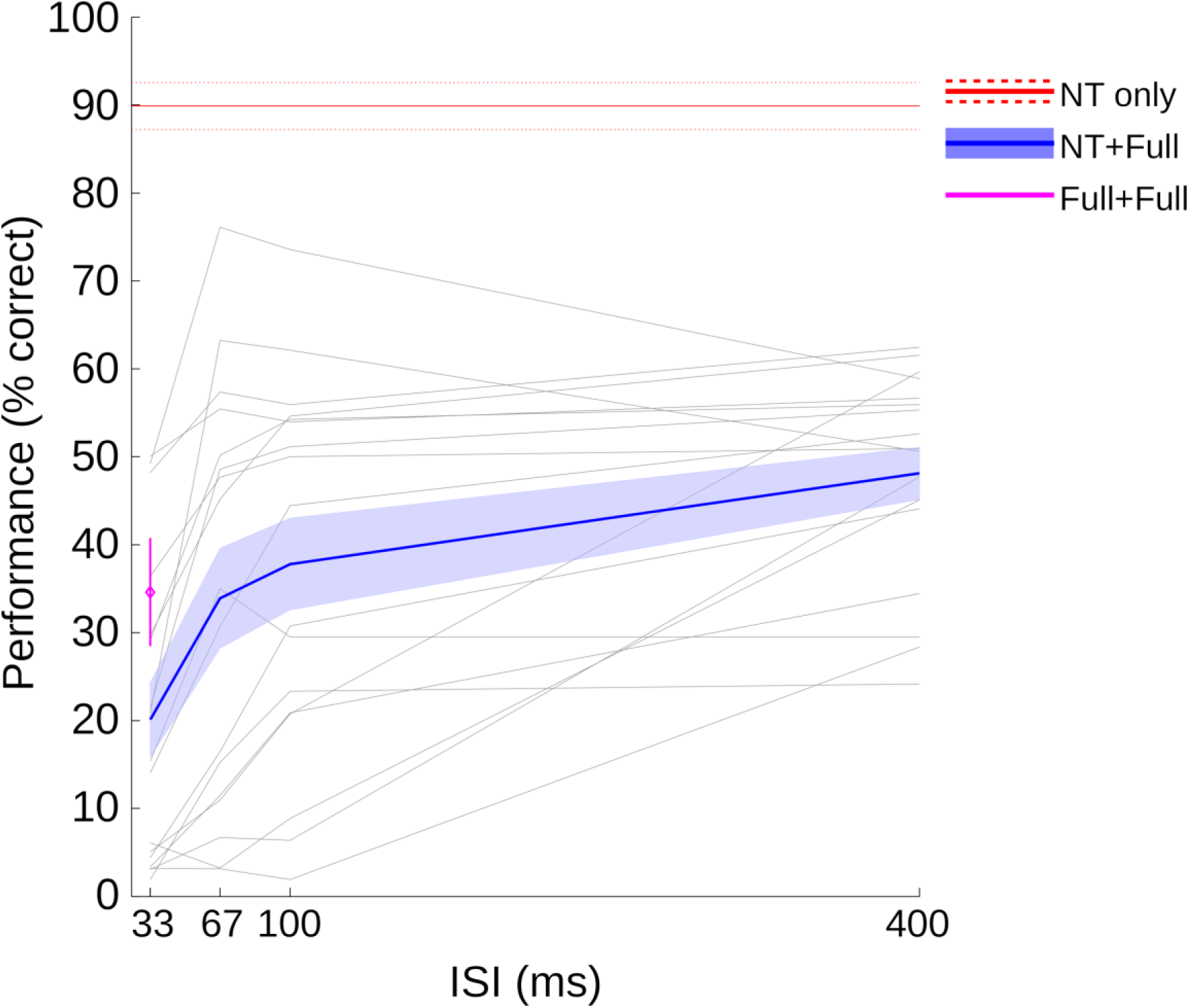
Percent correct responses in enumerating the number of flashes (only including trials in which the flash was correctly localized in spatial location). Blue lines indicate performance in the main experimental condition, in which one near-threshold flash and one full contrast stimulus were presented (NT + Full). The solid line shows the mean and the shaded area indicates the standard error of the mean across subjects. The red line shows performance in the near threshold (NT)-only condition (mean and s.e.m. across subjects). The magenta symbol, at 33ms, shows performance for two full pulses separated by a 33ms ISI (mean and s.e.m. across subjects). The faint gray lines represent performance of individual subjects in the NT + Full condition.

The main focus of the analysis was on the improvement in performance in the main experimental condition (NT + Full), as a function of ISI (Figure 4, blue line). As expected, subjects improved their performance in the NT + Full condition when the temporal separation between flashes was increased. This improvement leveled off between 100 and 400ms ISI, on average. Some subjects improved to levels near their individual performance maximum already by 67ms, while other subjects appeared to require longer ISIs to reach maximum performance (individual participant data shown as individual grey lines in Figure 4).

### MEG Data

The analysis of the MEG data was focused on three main aspects of the time series data. First, we analyzed stimulus-evoked (ERF) signatures for their relation to behavioral performance. Second, we determined the individual peak alpha frequency (PAF) for each participant and tested whether this was correlated with behavioral performance. Finally, we measured the trial-by-trial instantaneous alpha frequency (IAF) to see whether trials in which the alpha peak frequency was higher led to better performance in the two-flash task.

#### Event Related Fields

We first investigated confirmed that the NT and full strength stimuli were processed differently, with a greater response to the full strength flash. As expected, posterior/occipital and parietal sensor clusters responded based on stimulus strength, with weakest responses to the near-threshold stimulus and strongest response to two full contrast stimuli, with the ERF for the combination of a near-threshold and a full contrast stimulus falling somewhere in between, often a reduced version of the Full + Full condition. Locally averaged magnetometer ERFs (Figure 5) as well as general topographic time course (Figure 6, left) suggest the presence of spatially large, lateral dipoles involved in visual processing. Gradiometers show strong earlier activations mostly in posterior (occipital) regions, while later activity appears to transition to more central locations (Figure 6, right). To further identify regions that were sensitive to the stimulation strength, the time course of the pooled NT + Full conditions was subtracted from the Full + Full condition (see Figure 7).

**Figure 5:**
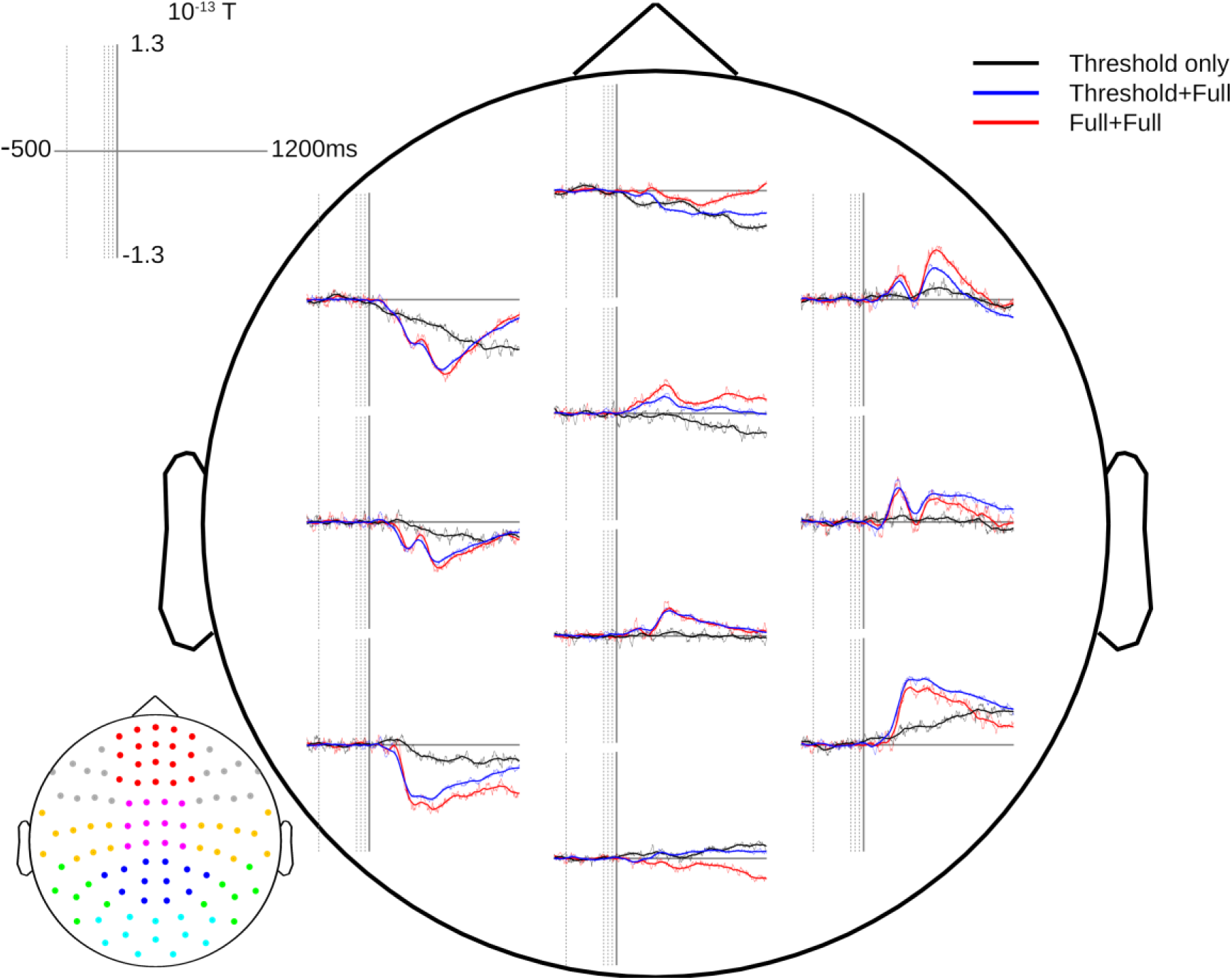
Topographic overview of averaged ERFs (magnetometers). For the purpose of illustration, NT + Full conditions (with ISIs of 33, 67, 100 and 400ms) have been averaged across ISI value for better signal-to-noise ratio. The lower left plot illustrates sensor averaging. The four faint vertical gray lines mark the temporal positions of the threshold stimuli in the NT + Full condition. Thin lines represent Maxfiltered data, thick lines have additionally been smoothed with a moving 100ms window (approximating 10Hz lowpass). Baseline and detrending was performed on [-500 0]ms interval.

**Figure 6:**
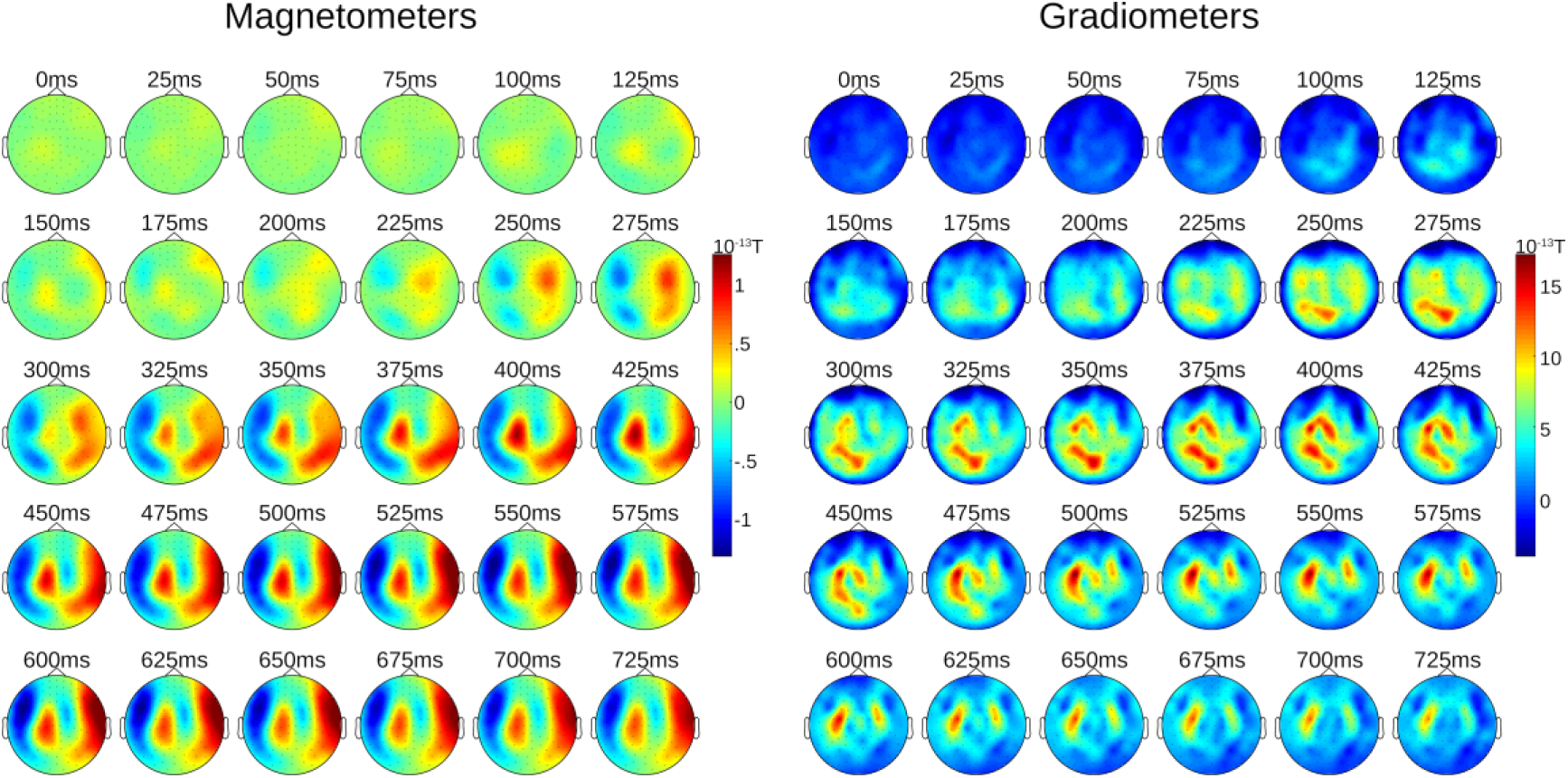
Topographic Time Course. Magnetometers and combined gradiometers, based on the averaged NT + Full conditions. Time values given indicate the central sample of a 25ms window. The average of [-50ms 0ms] has been subtracted for baseline normalization.

**Figure 7:**
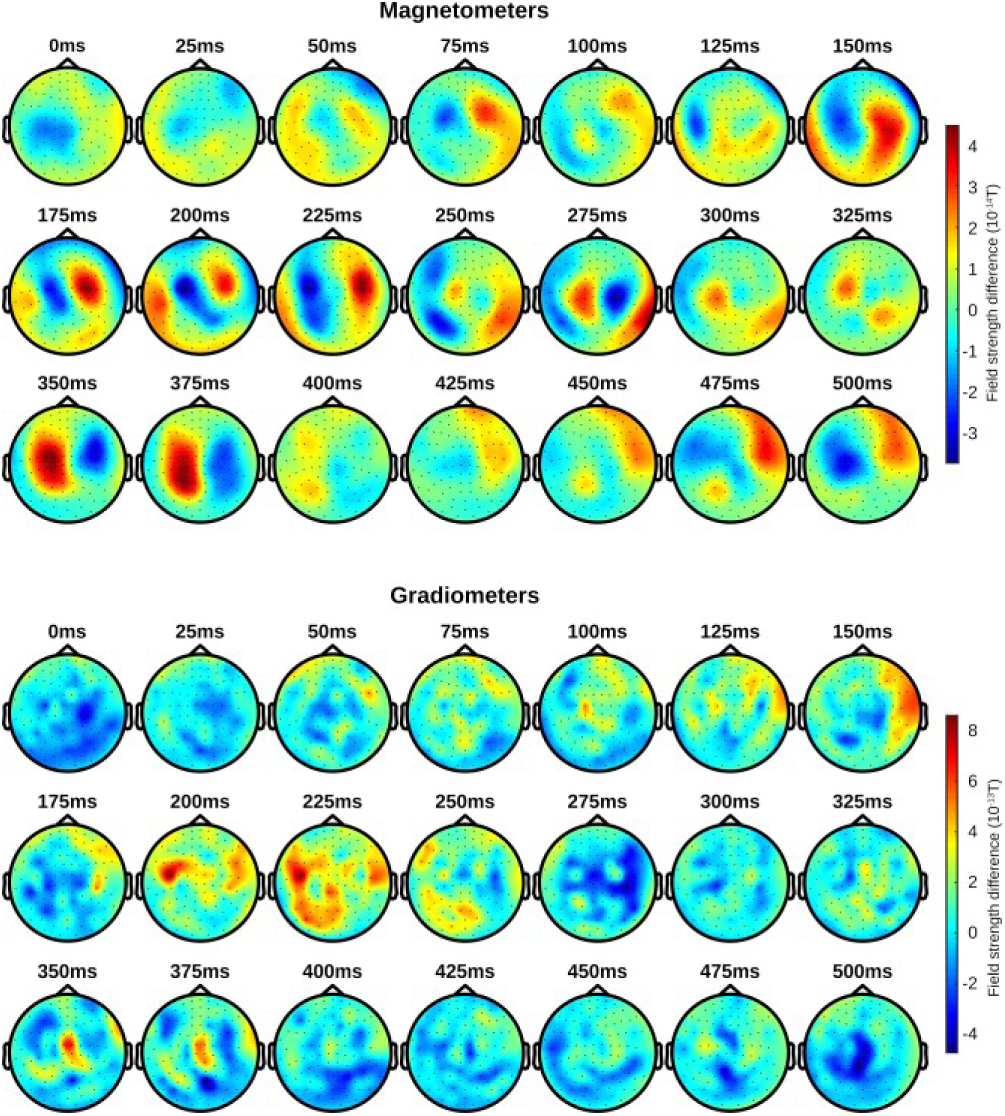
Topography of contrast (difference) between Full+Full and NT+Full conditions (Full+Full – NT+Full). The average of [-50ms 0ms] has been subtracted for baseline normalization. Top group: Magnetometers. Bottom group: Combined gradiometers.

**Figure 8:**
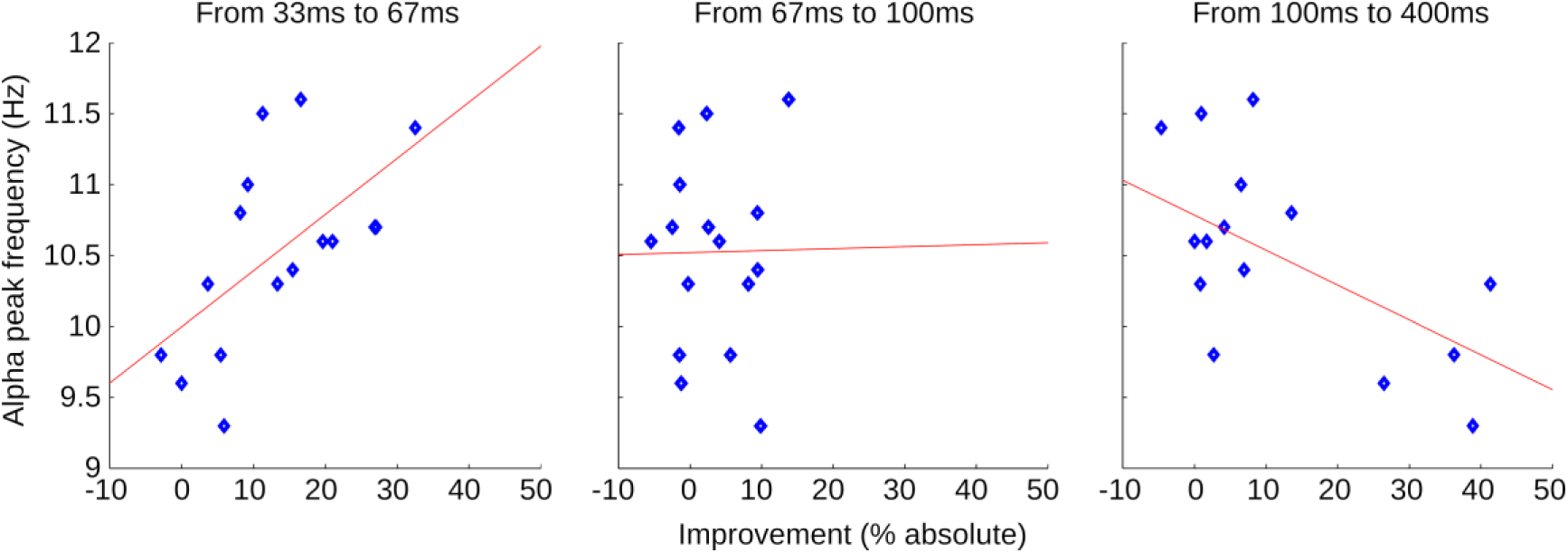
Peak Alpha Frequency vs. Behavioral Performance. Blue markers represent individual subjects; red lines are fitted to the data for better visualization. Results indicate a positive correlation between individual PAF and performance improvement in the early interval (33-67ms, Pearson’s R=0.60, p=0.014). No correlation was observed in the intermediate interval (67-100ms, R=0.01, p=0.966), whereas a negative correlation was found in the late interval (100-400ms, R=-0.60, p=0.013).

#### Correlation Analysis: Individual Peak Alpha Frequency

The behavioral data showed a common pattern: all subjects exhibited relatively poor performance at an ISI of 33ms, but also relatively high performance at 400ms ISI. The distribution of the improvement from the shortest to the longest timing varied between subjects; some subjects showed a strong increase in the earliest interval (33-67ms), while other participants showed the largest improvement in the latest interval (100-400ms).

Individual subject performance in this perceptual segregation task should be better for subjects with faster temporal resolution. Under the assumption that the individual resting state alpha peak frequency is an indicator of the individual speed of a subjects’ visual system and thereby of the ability to resolve brief temporal events (Cecere et al., 2015; Samaha and Postle, 2015; Wutz et al., 2018), we hypothesize that those subjects with higher individual peak alpha frequency (PAF) will improve their performance in earlier interstimulus intervals compared to subjects with slower PAF. Subjects with high individual PAF should therefore improve mostly in the first interval (33-67ms), subjects with low PAF should improve mostly in the last interval (100-400ms).

To test this hypothesis, we performed a correlation analysis between the individual resting state alpha peak frequencies of our subjects (see Figure 3) and their respective performance improvements between the 4 ISIs of the Threshold-Full condition (see the gray lines in Figure 4).

Results are shown in Figure 9. We found a significant positive correlation between the performance improvement and individual alpha peak frequency in the first interval (33-67ms, Pearson’s R=0.60, p=0.014). No correlation was observed in the intermediate interval (67-100ms, R=0.01, p=0.966), whereas a significant negative correlation was found in the late interval (100-400ms, R=-0.60, p=0.013). Note that reported p-values are uncorrected, but remain significant after Bonferroni-correction (N=3, threshold <=0.05).

**Figure 9:**
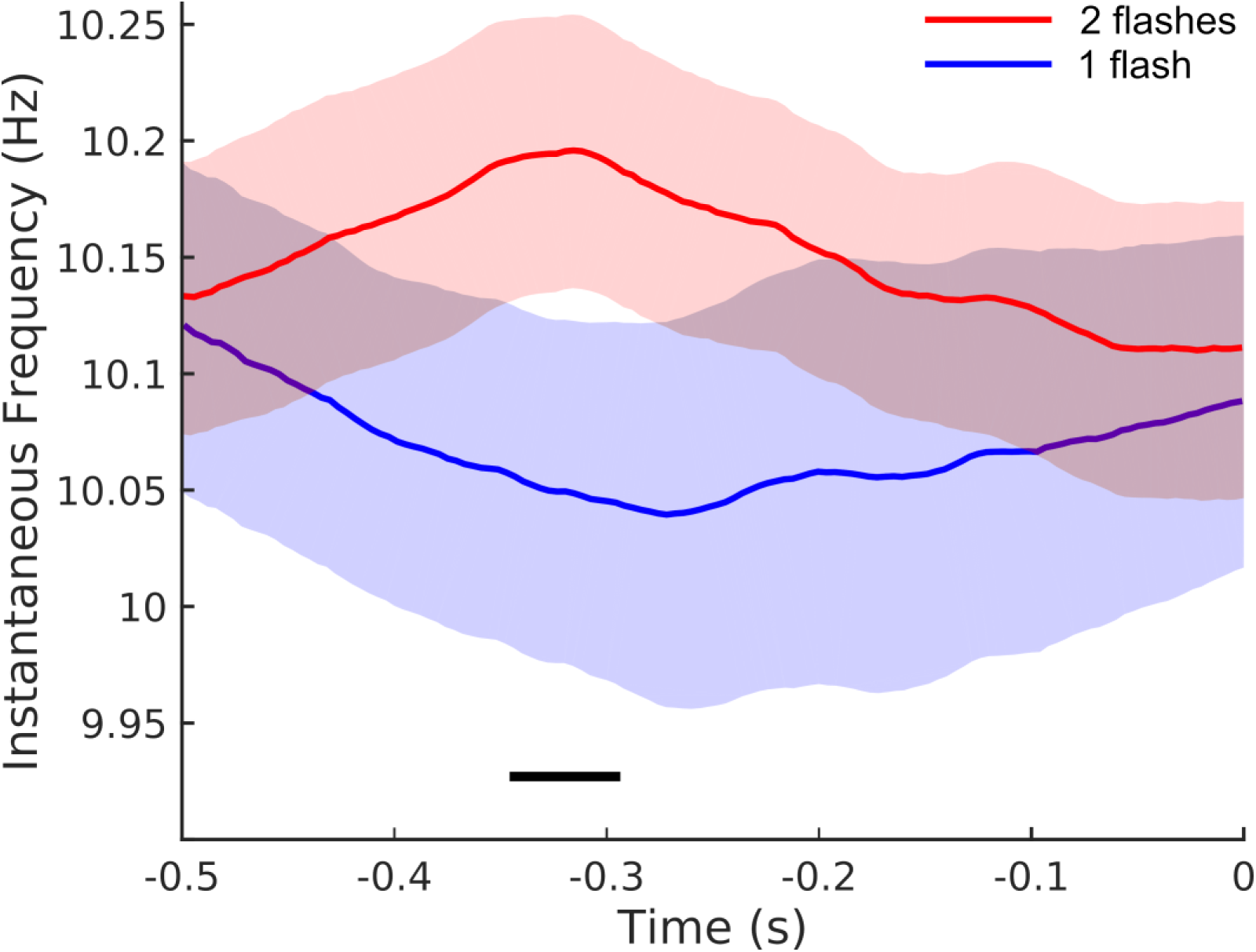
Pre-stimulus instantaneous alpha frequency. Within-subjects analysis comparing NT + Full trials in which subjects correctly reported two flashes (red) vs. those trials in which subjects reported only one flash (blue), mean across subjects and within-subject s.e.m.. The black line indicates the temporal location of a significant difference (cluster-corrected permutation test; p<0.05). The legend indicates the perceptual outcome as reported by the subjects, since all trials were in fact two flash presentations (NT + Full).

#### Correlation Analysis: Pre-Stimulus Instantaneous Alpha Frequency

As mentioned in the introduction, alpha frequency is generally considered to be a long-term stable feature (Salinsky et al., 1991). However, on a shorter time scale, peak oscillatory frequency (in the alpha band) varies within an individual during visual perception (Cohen, 2014; Samaha and Postle, 2015; Wutz et al., 2018). It has further been shown that just before stimulus onset, correctly discriminated (segregated) visual stimuli in a temporal fusion task on average exhibit slightly higher instantaneous alpha frequency (IAF) compared to incorrectly discriminated (fused) stimuli (Samaha and Postle, 2015; Wutz et al., 2018).

Consistent with this prediction, instantaneous alpha-band frequency in our subjects was higher on average for trials in which both flashes were reported, compared to those trials where only one flash was reported (Figure 9). Permutation testing revealed a significant difference between correctly and incorrectly reported stimuli over an interval from 344 to 296ms before stimulus onset (p= 0.013, cluster corrected).

## Discussion

Using a modified version of the two-flash fusion task, in which we also included pairs made up of a near-threshold (NT) stimulus and a suprathreshold stimulus with a varied inter-stimulus interval (based on (Wühle et al., 2010, 2011)), we investigated the link between alpha frequency and the processing of two rapidly presented stimuli. Consistent with the aim of our experimental design, the initial NT stimulus did not generate a strong event-related signal, meaning that the NT+full paradigm can indeed serve as a useful tool for investigating temporal integration without the risk of the first stimulus causing a strong phase-reset (Brandt, 1997; Wutz et al., 2014b) or artificially generating an “oscillation-like” activity (VanRullen and Macdonald, 2012). Further studies with this type of stimulus, using a more parametric mapping of ISIs, could be used (as in the Wühle and colleagues studies) to map the effective temporal integration window of different areas in MEG source space.

Our first hypothesis was that individual differences in the peak alpha frequency (PAF), measured during a separate resting-state period, would be related to individual differences in temporal resolution. We used the degree to which performance increased rapidly as a function of ISI (showing high temporal resolution) as a proxy measure for the speed of visual sampling. Consistent with our hypothesis, participants with a faster PAF showed the most rapid improvement in performance: they improved as ISI increased up to around 100ms, but then no longer benefitted from the longer ISI. Conversely, those with slower PAF did not benefit much from the initial increase in ISI but did improve more when ISI was over 100ms. This pattern of results provides further evidence for a link between resting state PAF and visual temporal resolution.

The second main finding was that the instantaneous measure of peak alpha frequency (IAF), estimated on each trial for the pre-stimulus period, was indeed higher/faster when participants correctly reported 2 flashes on that trial compared to when they only reported one flash. This provides a replication of a previous finding with the more traditional two-flash fusion task (Samaha and Postle, 2015), but without the possible confound of the first flash resetting the alpha phase. It was also interesting to show this effect in the current study given the other finding, described above, linking performance to individual differences. In terms of the question of whether the link between visual temporal resolution and alpha rhythms reflects a “trait” or a “state”, our findings suggest that both may be the case.

The overall pattern of results found here provides converging evidence for the claim that the “frame rate” of object processing in vision is linked to a specific visual temporal integration window that depends on the temporal sampling rate, at around 10Hz, of visual object processing. This is not to say that every aspect of visual (or multisensory) processing operates at this same frame rate (see, for example: (Ronconi et al., 2017)). Some forms of flicker, for example, are visible at faster rates (Landis, 1953; Brindley et al., 1966; Campos and Bedell, 1978; Capilla and Aguilar, 1993). One possible interpretation of the current set of findings is that rapid individuation of objects as distinct entities in space and time involves this 10Hz rhythm, while processing of more complex objects and events may involve slower rhythms, in the theta range (Drewes et al., 2015; Wutz et al., 2016; Zhu et al., 2016; Ronconi et al., 2017). Moreover, there is converging evidence that selective attention may also alternate, in terms of its function or spatial locus, in the theta to low-alpha range (Landau and Fries, 2012; Fiebelkorn et al., 2013; VanRullen, 2013; Song et al., 2014; Dugué et al., 2015).

In conclusion, our results addressed the question of whether the individual peak alpha frequency (PAF), linked to a visual sampling rhythm, is an individual trait (PAF) or state-dependent and subject to change via top-down control (Samaha and Postle, 2015; Wutz et al., 2018) or other factors. The term “instantaneous alpha frequency” (IAF) suggests variability over time. Here, we found that overall performance was indeed linked to an individual trait (PAF), but also that variations in the IAF predicted performance on individual trials. In other words, peak alpha may be both a trait and a state.

## Acknowledgments

We would like to thank Poppy Sharp for her assistance with data collection, and Andreas Wutz for his help with data analysis. JD, EM and DM were supported by a European Research Council (ERC) grant (Grant Agreement No. 313658) and High-level Foreign Expert Grant (GDT20155300084). WZ was supported by the National Natural Science Foundation of China (61263042, 61563056).

## References

Aurlien H, Gjerde IO, Aarseth JH, Eldøen G, Karlsen B, Skeidsvoll H, Gilhus NE (2004) EEG background activity described by a large computerized database. Clin Neurophysiol Off J Int Fed Clin Neurophysiol 115:665–673.

Brainard DH (1997) The Psychophysics Toolbox. Spat Vis 10:433–436.

Brandt ME (1997) Visual and auditory evoked phase resetting of the alpha EEG. Int J Psychophysiol 26:285–298.

Brindley GS, Du Croz JJ, Rushton WAH (1966) The flicker fusion frequency of the blue-sensitive mechanism of colour vision. J Physiol 183:497–500.

Campos EC, Bedell HE (1978) Critical flicker-fusion frequency as an indicator of human receptive field-like properties. Invest Ophthalmol Vis Sci 17:533–538.

Capilla A, Pazo-Alvarez P, Darriba A, Campo P, Gross J (2011) Steady-State Visual Evoked Potentials Can Be Explained by Temporal Superposition of Transient Event-Related Responses. PLoS ONE 6 Available at: https://www.ncbi.nlm.nih.gov/pmc/articles/PMC3022588/ [Accessed February 27, 2020].

Capilla P, Aguilar M (1993) Red-green flicker resolution as a function of heterochromatic luminous modulation. Ophthalmic Physiol Opt J Br Coll Ophthalmic Opt Optom 13:183–185.

Cecere R, Rees G, Romei V (2015) Individual differences in alpha frequency drive crossmodal illusory perception. Curr Biol CB 25:231–235.

Cohen MX (2014) Fluctuations in Oscillation Frequency Control Spike Timing and Coordinate Neural Networks. J Neurosci 34:8988–8998.

Dehaene S (2016) Temporal Oscillations in Human Perception. Psychol Sci Available at: https://journals.sagepub.com/doi/10.1111/j.1467-9280.1993.tb00273.x [Accessed February 27, 2020].

Doelling KB, Assaneo MF, Bevilacqua D, Pesaran B, Poeppel D (2019) An oscillator model better predicts cortical entrainment to music. Proc Natl Acad Sci 116:10113–10121.

Drewes J, Zhu W, Wutz A, Melcher D (2015) Dense sampling reveals behavioral oscillations in rapid visual categorization. Sci Rep 5:16290.

Dugué L, Marque P, VanRullen R (2015) Theta oscillations modulate attentional search performance periodically. J Cogn Neurosci 27:945–958.

Fiebelkorn IC, Saalmann YB, Kastner S (2013) Rhythmic Sampling within and between Objects despite Sustained Attention at a Cued Location. Curr Biol 23:2553–2558.

Gho M, Varela FJ (1988) A quantitative assessment of the dependency of the visual temporal frame upon the cortical rhythm. J Physiol (Paris) 83:95–101.

Haegens S, Cousijn H, Wallis G, Harrison PJ, Nobre AC (2014) Inter- and intra-individual variability in alpha peak frequency. NeuroImage 92:46–55.

Hasson U, Yang E, Vallines I, Heeger DJ, Rubin N (2008) A Hierarchy of Temporal Receptive Windows in Human Cortex. J Neurosci 28:2539–2550.

Klimesch W (1999) EEG alpha and theta oscillations reflect cognitive and memory performance: a review and analysis. Brain Res Rev 29:169–195.

Lakatos P, Gross J, Thut G (2019) A New Unifying Account of the Roles of Neuronal Entrainment. Curr Biol 29:R890–R905.

Landau AN, Fries P (2012) Attention samples stimuli rhythmically. Curr Biol CB 22:1000–1004.

Landis C (1953) An Annotated Bibliography of Flicker Fusion Phenomena Covering the Period 1740-1952. National Academies.

Oostenveld R, Fries P, Maris E, Schoffelen J-M (2011) FieldTrip: Open Source Software for Advanced Analysis of MEG, EEG, and Invasive Electrophysiological Data. Comput Intell Neurosci Available at: https://www.hindawi.com/journals/cin/2011/156869/ [Accessed October 14, 2019].

Oosterhof NN, Connolly AC, Haxby JV (2016) CoSMoMVPA: Multi-Modal Multivariate Pattern Analysis of Neuroimaging Data in Matlab/GNU Octave. Front Neuroinformatics 10 Available at: https://www.ncbi.nlm.nih.gov/pmc/articles/PMC4956688/ [Accessed April 7, 2020].

Pelli DG (1997) The VideoToolbox software for visual psychophysics: transforming numbers into movies. Spat Vis 10:437–442.

Pöppel E (1997) A hierarchical model of temporal perception. Trends Cogn Sci 1:56–61.

Pöppel E (2009) Pre-semantically defined temporal windows for cognitive processing. Philos Trans R Soc B Biol Sci 364:1887–1896.

Ronconi L, Melcher D (2017) The Role of Oscillatory Phase in Determining the Temporal Organization of Perception: Evidence from Sensory Entrainment. J Neurosci Off J Soc Neurosci 37:10636–10644.

Ronconi L, Oosterhof NN, Bonmassar C, Melcher D (2017) Multiple oscillatory rhythms determine the temporal organization of perception. Proc Natl Acad Sci U S A 114:13435–13440.

Salinsky MC, Oken BS, Morehead L (1991) Test-retest reliability in EEG frequency analysis. Electroencephalogr Clin Neurophysiol 79:382–392.

Samaha J, Postle BR (2015) The Speed of Alpha-Band Oscillations Predicts the Temporal Resolution of Visual Perception. Curr Biol 25:2985–2990.

Song K, Meng M, Chen L, Zhou K, Luo H (2014) Behavioral Oscillations in Attention: Rhythmic α Pulses Mediated through θ Band. J Neurosci 34:4837–4844.

Taulu S, Kajola M (2005) Presentation of electromagnetic multichannel data: The signal space separation method. J Appl Phys 97:124905.

Taulu S, Simola J, Kajola M (2005) Applications of the signal space separation method. IEEE Trans Signal Process 53:3359–3372.

VanRullen R (2013) Visual Attention: A Rhythmic Process? Curr Biol 23:R1110–R1112.

VanRullen R, Koch C (2003) Is perception discrete or continuous? Trends Cogn Sci 7:207–213.

VanRullen R, Macdonald JSP (2012) Perceptual Echoes at 10 Hz in the Human Brain. Curr Biol 22:995–999.

Varela FJ, Toro A, John ER, Schwartz EL (1981) Perceptual framing and cortical alpha rhythm. Neuropsychologia 19:675–686.

Watson AB, Pelli DG (1983) QUEST: a Bayesian adaptive psychometric method. Percept Psychophys 33:113–120.

Wühle A, Mertiens L, Rüter J, Ostwald D, Braun C (2010) Cortical processing of near-threshold tactile stimuli: an MEG study. Psychophysiology 47:523–534.

Wühle A, Preissl H, Braun C (2011) Cortical processing of near-threshold tactile stimuli in a paired-stimulus paradigm--an MEG study. Eur J Neurosci 34:641–651.

Wutz A, Melcher D, Samaha J (2018) Frequency modulation of neural oscillations according to visual task demands. Proc Natl Acad Sci 115:1346–1351.

Wutz A, Muschter E, van Koningsbruggen M, Melcher D (2014a) Saccades reset temporal integration windows. J Vis 14:584–584.

Wutz A, Muschter E, van Koningsbruggen MG, Weisz N, Melcher D (2016) Temporal Integration Windows in Neural Processing and Perception Aligned to Saccadic Eye Movements. Curr Biol 26:1659–1668.

Wutz A, Weisz N, Braun C, Melcher D (2014b) Temporal Windows in Visual Processing: “Prestimulus Brain State” and “Poststimulus Phase Reset” Segregate Visual Transients on Different Temporal Scales. J Neurosci 34:1554–1565.

Yamashiro K, Inui K, Otsuru N, Urakawa T, Kakigi R (2011) Temporal window of integration in the somatosensory modality: An MEG study. Clin Neurophysiol 122:2276–2281.

Zhu W, Drewes J, Melcher D (2016) Time for Awareness: The Influence of Temporal Properties of the Mask on Continuous Flash Suppression Effectiveness. PLOS ONE 11:e0159206.

Zoefel B, ten Oever S, Sack AT (2018) The Involvement of Endogenous Neural Oscillations in the Processing of Rhythmic Input: More Than a Regular Repetition of Evoked Neural Responses. Front Neurosci 12 Available at: https://www.frontiersin.org/articles/10.3389/fnins.2018.00095/full [Accessed February 27, 2020].

